# PD-L1 expression in 522 selected sarcomas with subset analysis of recurrent or metastatic matched samples and association with tumour-infiltrating lymphocytes

**DOI:** 10.1101/757625

**Authors:** Ana Cristina Vargas, Fiona M Maclean, Loretta Sioson, Dinh Tran, Fiona Bonar, Annabelle Mahar, Alison L. Cheah, Peter Russell, Peter Grimison, Louise Richardson, Anthony J Gill

## Abstract

We assessed the frequency of programmed death-ligand 1 (PD-L1) expression by immunohistochemistry (IHC) in a cohort of 522 sarcomas from 457 patients, incuding a subset of 46 patients with 63 matched samples from local recurrence or metastases with primary tumours and/or metachronous metastases. We also investigated the correlation of PD-L1 with the presence and degree of tumour-infiltrating lymphocytes (TILs) in a subset of cases. IHC was performed using the PD-L1 SP263 companion kit (VENTANA) on tissue microarrays from an archival cohort. Evaluation of PD-L1 and TILs was performed on full sections for a subset of 23 cases. Fisher’s exact and Mann Whitney test were used to establish significance (P <0.05). PD-L1 positive expression (≥1%) was identified in 31% of undifferentiated pleomorphic sarcomas, 29% of angiosarcomas, 26% of rhabdomyosarcomas, 18% of myxofibrosarcomas, 11% of leiomyosarcomas and 10% of dedifferentiated liposarcomas. Negative expression was present in all atypical lipomatous tumous/well-differentiated lipoasarcomas, myxoid liposarcomas, synovial sarcomas, pleomorphic liposarcomas, and Ewing sarcomas. PD-L1 IHC was concordant in 81% (38 of 47) of matched/paired samples. PD-L1 IHC was discordant in 19% (9 of 47 matched/paired samples), displaying differences in the proportion of cells expressing PD-L1 amongst paired samples with the percentage of PD-L1-positive cells increasing in the metastatic/recurrent site compared to the primary in 6 of 9 cases (67%). Significant correlation between PD-L1 expression and the degree of TILs was exclusively identified in the general cohort of leiomyosarcomas, but not in other sarcoma subtypes or in metastatic/recurrent samples. We conclude that the prevalence of PD-L1 expression in selected sarcomas is variable and likely to be clone dependent. Importantly, we demonstrated that PD-L1 can objectively increase in a small proportion of metastases/recurrent sarcomas, offering the potential of treatment benefit to immune checkpoint inhibitors in this metastatic setting.

## Introduction

The PD-1/PD-L1 axis (Programmed death 1 or CD274/ programmed death-ligand 1) plays a crucial role in immune surveillance. PD-1 is a transmembrane protein expressed on activated T and B cells and binds to its ligands, PD-L1 or PD-L2, which are variably expressed in immune (T and B cells, dendritic cells, mast cells) and non-immune (e.g. endothelial) cells including tumour cells. The PD-1/PD-L1 axis acts as an immune checkpoint by inhibiting T-cell function leading to tumour immune escape (Reviewed in(1, 2)). Therapeutic blockage of the PD-1/PD-L1 receptor–ligand can result in durable clinical responses in lung and bladder cancer (2–6) but their benefit in various sarcoma subtypes remains unknown, with no phase 3 studies yet published.

The frequency of PD-L1 expression in sarcomas reported in the literature is highly variable with incidences ranging from 0% to 65%(5, 7–19). In their analysis, some of these series have combined statistical analysis of the two molecularly characteristic groups of sarcomas, that is those associated with recurrent specific genetic events (e.g. translocations or amplification) known to drive tumorogenesis (e.g. t(X;18)(p11.2; q11.2) in synovial sarcoma) and sarcomas with complex karyotype (e.g. myxofibrosarcoma), which lack detectable recurrent gene alterations, as a single entity (20–23). Such distinction is fundamental as sarcomas across the different groups and within the same group show distinct clinical behaviour.

The predictive and prognostic significance of PD-L1 expression in sarcomas has also been reviewed including its correlation with PD-1 expression, and the presence and degree of tumour-infiltrating lymphocytes (TILs) (5, 7–19).

The aims of our study were threefold. To review the incidence of PD-L1 expression in a cohort of 522 selected bone and soft tissue sarcomas from 457 patients with the commercially available and widely used PD-L1 companion kit SP263 (Ventana)(24). This immunohistochemical assay in other tumour types (i.e. lung and urothelial) can identify patients more likely to benefit from treatment with anti-PD-[L]1 immunotherapy such as durvalumab, pembrolizumab and nivolumab. Only few studies using the clone SP263 have been published in sarcoma patients (10, 18). A second aim was to review the level of concordance in PD-L1 expression in matched sarcoma samples from primary tumour and its recurrence and/or metastasis (or metachronous recurrent/metastatic episodes). The purpose was to identify whether changes in PD-L1 expression occur during the evolution of more aggressive sarcomas as if so, these may impact treatment of refractory advanced disease. Lastly, we correlated PD-L1 expression with the presence and degree of TILs identified on H&E sections for a subset of cases. The significance of TILs identified on routine H&E sections in sarcomas has not been widely investigated.

## Materials and Methods

This study was approved by the Northern Sydney Local Health District (NSLHD) Human Research Ethics Committee (HREC) reference 1312-417M. A retrospecitve database search was performed of the archives of Douglass Hanly Moir (DHM) Pathology laboratory in a 10-year (2008-2018) period of time to identify a cohort of selected bone and soft tissue sarcomas. These were diagnosed by pathologists with sarcoma expertise following standardised diagnostic criteria(20), which included the used of FISH probes (Break-apart: SS18, EWSR1, FOXO1, DDIT3, FUS; and enumeration: MDM2) when required for tumour classification. As this is a de-identified cohort, follow-up is not available. Ewing sarcomas were the only primary bone tumours included in this cohort, all of which had an associated soft tissue component not requiring prior decalcification. Only formalin-fixed paraffin-embedded (FFPE) tissue blocks with adequate viable tumour from resection specimens were selected for construction of tissue microarrays (TMAs). Replicate tissue microarray (TMA) cores were taken from each sample (2×1 mm cores). For PD-L1 IHC, paraffin blocks were sectioned at 4 μm on to Superfrost Plus slides and stained with the Ventana PD-L1 SP263 rabbit monoclonal primary antibody (Assay v1.00.0001), using the VENTANA OptiView DAB IHC Detection Kit and the BenchMark ULTRA platform. IHC was performed on TMAs for the entire cohort and on full sections for a subset of cases (n=23). Positive (normal placenta as described (25) and negative controls were included in each run. Positive PD-L1 expression in the tumour cells was scored semiquantitatively as 0-3+, based on weak, moderate and strong membranous staining, respectively and the percentage of tumour cells expressing the protein was recorded. Following standard guidelines as per lung for the clone SP263 (24), PD-L1 expression was regarded as positive if complete or incomplete membranous expression was present in ≥1% of the tumour cells regardless the intensity. PD-L1 expression on immune cells (lymphocytes and/or plasma cells) was also recorded when identified. Although we aimed to score PD-L1 on 100 viable tumour cells, this was not achievable for sarcomas with low cellularity on TMAs. Nevertheless, standardised criteria for PD-L1 scoring, including the minimal number of cells for PD-L1 expression, in the context of sarcoma samples has not been universally established.

Evaluation of TILs was performed on 2 to 3 representative full H&E sections (4 microns in thickness) per each case only for the sarcoma subtypes with the largest number of cases (UPS, MFS and LMS) and for the metastatic/recurrent samples (Met/Rec cohort: n=306). TILs were scored following the recommendations by the International TILs Working Group for breast cancer(31). Briefly, the intratumoural area occupied by stromal TILs was expressed as a percentage. TILs were assessed in areas away from necrosis. Tertiary lymphoid structures and other inflammatory cells (i.e. neutrophils and histiocytes) were excluded. The level of TILS was regarded as nil (0%), minimal (1-10%), intermediate (>10% and <50%) and high (>50%). These cut-offs were selected following the groups A-C of Salgado et al (26–27), with the exception that for this study, absent TILs (0%) was differentially recorded from mild TILs (1-10%). A limitation of applying this system to sarcomas is that, contrary to breast cancer, the tumour boundary in a proportion of sarcoma subtypes is not well defined.

For statistical analysis the Fisher exact test for a 2 × 2 contingency table and the Mann Whitney test were used to establish significant correlations between PD-L1 and TILs with significance determined at P value: <0.05. An online calculator was used for the analysis: https://www.socscistatistics.com/tests/fisher/default2.aspx and https://www.socscistatistics.com/tests/mannwhitney/.

## RESULTS

### Overall cohort

The entire cohort assessed for PD-L1 IHC on TMAs comprised 522 samples from 457 patients. These included primary uterine and soft tissue leiomyosarcomas (LMS n=116), atypical lipomatous tumours/well- differentiated liposarcomas (ALT/WD-LPS, n=95), myxofibrosarcomas (MFSs, n=97), undifferentiated pleomorphic sarcomas (UPS, n=67), dedifferentiated liposarcomas (DD-LPS, n=37), myxoid liposarcomas (ML, n=27), rhabdomyosarcomas (RMS, n=24: 9 pleomorphic, 7 embryonal, 4 alveolar and 3 spindle cell/sclerosing), synovial sarcomas (SS, n=17), angiosarcomas (n=17), Ewing sarcomas (ES, n=14) and pleomorphic liposarcomas (n=11).From this cohort, 63 samples were derived from recurrent/metastatic specimens (plus residual disease for a few of these cases, **Table I**) referred to as Rec/met cohort, from 46 patients.

**Table I Title.** Matched sarcoma samples from 46 pateints (Met/Rec cohort) with results for PD-L1 IHC and tumour-infitlrating lymphocytes (TILs).

| Case No. | Label ID. | Patient's age | Diagnosis | Site | Sample type | Year of Dx. | PD-L1 IHC % | TILs |
| --- | --- | --- | --- | --- | --- | --- | --- | --- |
| 1 | MFS-46<br>MFS-09 | 80 | Myxofibrosarcoma, Gde 3, 45mm | Calf<br>Calf | Primary<br>Rec | 14<br>16 | 0<br>0 | High<br>No TILs |
| 2 | MFS-92<br>MFS-71<br>MFS-72 | 65 | Myxofibrosarcoma, Gde 3, 115 mm | Buttock<br>Thigh<br>Thigh | Primary<br>Rec 1<br>Rec 2 | 11<br>12<br>12 | *5%<br>1%<br>*70% | No TILs<br>No TILs<br>No TILs |
| 3 | MFS-77<br>MFS-70<br>MFS-31<br>MFS-29<br>MFS-38 | 76 | Myxofibrosarcoma, Gde 3, 38mm, post-irradiation | Tibia<br>Tibia<br>Leg<br>Shin<br>Shin | Primary<br>Rec 1<br>Rec 2<br>Residual<br>Rec 3 | 12<br>13<br>13<br>13<br>14 | *1%<br>5%<br>*5%<br>1%<br>*10% | No TILs<br>No TILs<br>No TILs<br>Moderate<br>No TILs |
| 4 | MFS-65<br>MFS-64<br>MFS-52 | 48 | Myxofibrosarcoma, Gde 3, 35 mm | Thigh<br>Thigh<br>Thigh | Primary<br>Residual<br>Rec | 16<br>16<br>17 | 0<br>0<br>0 | No TILs<br>No TILs<br>No TILs |
| 5 | MFS-26<br>MFS-74 | 65 | Myxofibrosarcoma, Gde 3 20mm, post-irradiation | Ulna<br>Forearm | Rec 1<br>Rec 2 | 10<br>12 | *1%<br>*<1% | Moderate, het<br>No TILs |
| 6 | MFS-17<br>MFS-22 | 68 | Myxofibrosarcoma, Gde 3, 22m | Arm<br>Arm | Rec 1<br>Rec 2 | 09<br>10 | 0<br>0 | Moderate<br>High |
| 7 | MFS-16<br>MFS-35<br>MFS-33<br>MFS-11 | 79 | Myxofibrosarcoma Gde 2, 40 mm; post-irradiation | Calf<br>Calf<br>Calf<br>Calf | Rec 1<br>Rec 2<br>Residual<br>Rec 3 | 09<br>13<br>13<br>14 | *5%<br>0<br>1%<br>*1% | Moderate, het<br>No TILs<br>No TILs<br>No TILs |
| 8 | MFS-43<br>MFS-60 | 90 | Myxofibrosarcoma, Gde 3, 54 mm | Thigh<br>Thigh | Rec 1<br>Rec 2 | 15<br>16 | *1%<br>*5% | Milderate, het<br>No TILs |
| 9 | MFS-84<br>MFS-47 | 60 | Myxofibrosarcoma, grade 3, 85 mm | Buttock<br>Brain | Primary<br>Met | 13<br>14 | *1%<br>*1% | No TILs<br>No TILs |
| 10 | MFS-80<br>MFS-02A | 67 | Myxofibrosarcoma, Gde 2, 130mm | Forearm<br>Forearm | Primary<br>Rec | 13<br>15 | 0<br>0 | Mild<br>Mild |
| 11 | MFS-69<br>MFS-42<br>MFS-63 | 83 | Myxofibrosarcoma, Gde 1 - 45mm | Leg<br>Leg<br>Leg | Rec 1<br>Rec 2<br>Rec 3 | 13<br>15<br>16 | 0<br>0<br>0 | No TILs<br>No TILs<br>No TILs |
| 12 | MFS-76<br>MFS-07 | 65 | Myxofibrosarcoma, Gde 2, 33m | Thigh<br>Thigh | Primary<br>Residual | 11<br>11 | 0<br>0 | No TILs<br>No TILs |
| 13 | MFS-85<br>MFS-90 | 65 | Myxofibrosarcoma, Gde 2, 20mm | Knee<br>Knee | Primary<br>Rec | 11<br>11 | 0<br>0 | No TILs<br>No TILs |
| 14 | MFS-61<br>MFS-62 | 64 | Myxofibrosarcoma, Gde 2, 60 mm | Arm<br>Arm | Primary<br>Residual | 16<br>16 | 0<br>0 | Moderate, het<br>Moderate |
| 15 | MFS-41<br>MFS-48<br>MFS-45 | 93 | Myxofibrosarcoma, Gde 2, 155 mm | Leg<br>Leg<br>Leg | Primary<br>Residual<br>Rec | 14<br>14<br>14 | 0<br>0<br>0 | No TILs<br>No TILs<br>No TILs |
| 16 | MFS-55<br>MFS-56 | 84 | Myxofibrosarcoma, Gde 2, 25mm | Back<br>Back | Primary<br>Residual | 16<br>16 | 0<br>0 | Mild<br>Mild |
| 17 | MFS-73<br>MPNST1 | 68 | Myxofibrosarcoma, Gde 3, 90 mm | Arm<br>Arm | Primary<br>Rec | 12<br>13 | 0<br>0 | No TILs<br>No TILs |
| 18 | UPS-17<br>UPS-15<br>UPS-52 | 66 | UPS, Gde 3, 18 mm | External canal<br>External canal<br>Parotid | Primary<br>Residual<br>Rec | 16<br>16<br>17 | 0<br>0<br>0 | No TILs<br>Moderate, het<br>No TILs |
| 19 | UPS-04<br>UPS-14 | 40 | UPS, Gde 3, 18 mm | Lung<br>Lung | Mets 1<br>Mets 2 | 15<br>16 | 0<br>0 | High, het<br>No TILs |
| 20 | UPS-40<br>UPS-49 | 77 | UPS grade 3, 40mm | Buttock<br>Buttock | Primary<br>Rec | 14<br>15 | 0<br>0 | No TILs<br>No TILs |
| 21 | UPS-62<br>UPS-61 | 74 | UPS, Gde 3 67 mm | Groin<br>Groin | Rec 1<br>Rec 2 | 13<br>13 | 100%<br>100% | No TILs<br>No TILs |
| 22 | UPS-44<br>UPS-16 | 61 | UPS Grade 3, 50 mm, postradiotherapy | Forearm<br>Forearm | Primary<br>Rec | 14<br>16 | 0<br>0 | High<br>High |

In the overall cohort, PD-L1 positive expression (≥1%) was identified in 31% UPSs (21/67), 29% angiosarcomas (5/17), 26% RMSs (6/24, most of which were of the pleomorphic subtype), 18% MFSs (17/97), 11% LMSs (13/116) and 10% DD-LPS (4/37). PD-L1 was negative (<1%) in all ALT/WDL (n=95), myxoid liposarcomas (n=27), synovial sarcomas (n=17), pleomorphic liposarcomas (n=11) and Ewing sarcomas (n=15). When identified, PD-L1 staining was limited to tumour cells with no definitive epression present in immune cells. It is possible that for cases with strong tumour expression, this obscured expression in the immune cells. As previoulsy described, PD-L1 expression was seen in macrophages and endothelial cells but this was inconsistent in these cell types across the samples. Both complete and incomplete membranous expression pattern was identified and the intensity of the stain was highly variable (**Figure 1A-E**). Protein expression was overall concordant in replicate cores (**Figure 1F**).

**Figure 1 legend.**
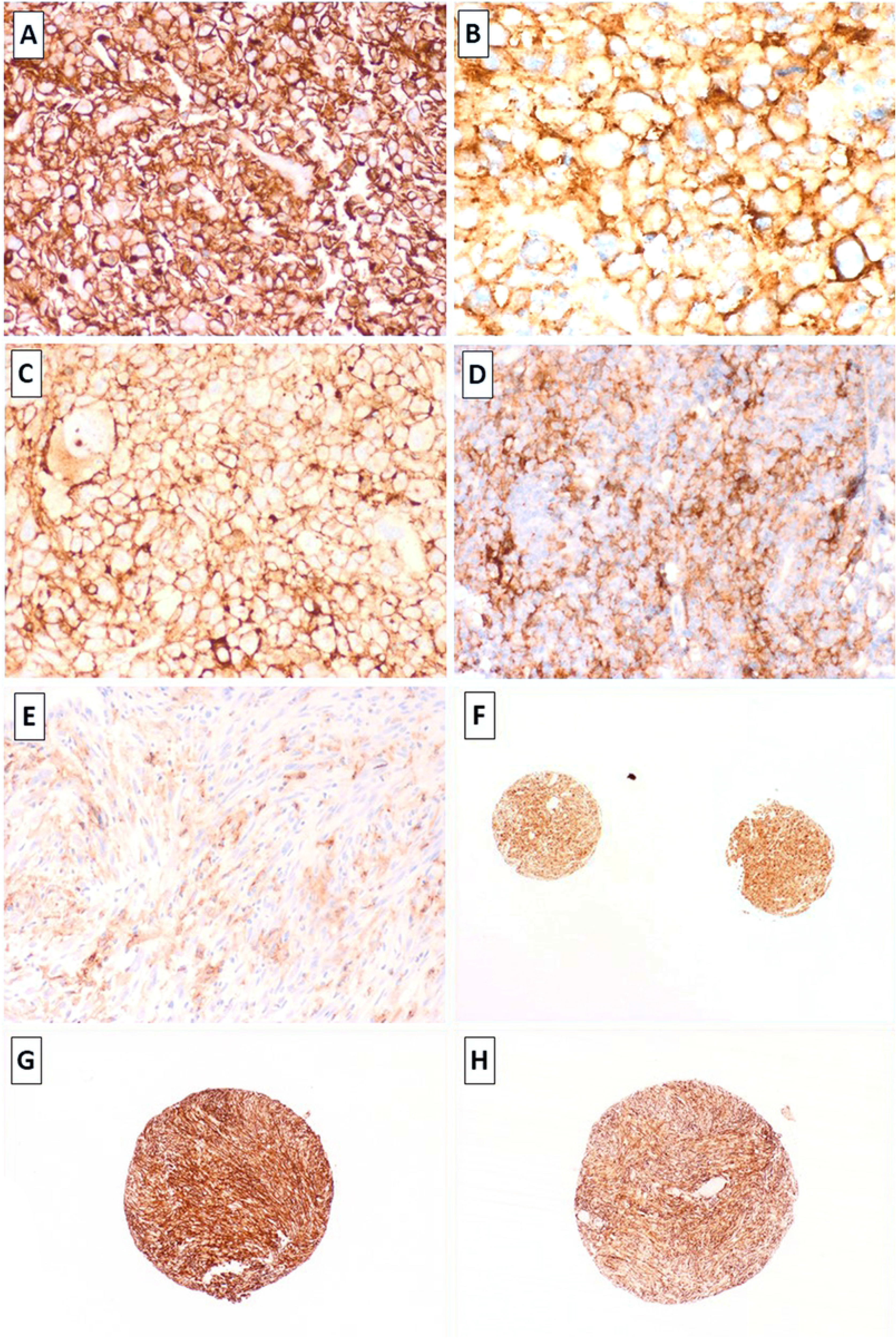
Strong membranous expression for PD-L1 idenified in leiomyosarcoma (LM-75), undifferentiated pleomorphic sarcomas (B,C: UPS-67 & UPS-11, respectively) and metastatic angiosarcoma (D, AS-2). Focal heterogenous expression for PD-L1 in leiomyosarcoma (E, LM99). Concordant expression for PD-L1 in replicate cores was identified (e.g. 1F; UPS-21). Primary and recurrent UPS (UPS-61 and UPS-62: G & H) demonstrating concordant strong expression of PD-L1 in 100% of the tumour cells in both, the primary lesion and its recurence (TMA cores). Images taken at 2x – 40x magnification and stained with the PD-L1 SP263 clone, Ventana.

### Rec / Met cohort

This includes paired samples from the following types: MFS (n=17), UPS (n=5), LMS (n=11), RMS (n=2), ML (n=2), pleomorphic LPS (n=2), angiosarcomas (n=3), SS (n=2), ES (n=1) and DD-LPS (n=1; **Table 1**). The median interval between acquisitions of paired lesions was variable (Range 0-6 years, **Table 1**). All paired samples including residual disease and those occurring <1 year apart were derived from independent surgical procedures (**Table 1**). These were included to assess for intra-tumour heterogeneity. Overall, PD-L1 was concordant amongst matched / paired samples for 79% of the cases when assessed on TMAs (37 of 46 patients). For the remaining 21% with discordant PD-L1 on TMAs, IHC was performed on full sections, which increased the level of concordance from 79% to 81%. A few of these cases were initially scored as negative on TMAs but on full sections, unequivocal PD-L1 stain was identified in >1% of the cells. Except for one, all cases lacked PD-L1 expression. Hence, these were negatively concordant (0% PD-L1 expression). The remaining case (UPS) displayed 100% expression concordantly in the primary and recurrent sample (**Figure 1G-H**). As this recurrence was identified soon after the resection of the primary tumour, it may represent residual disease.

**Table I Legend.** Results of PD-L1 IHC are displayed as scored on tissue microarrays (TMAs) when fully concordant. Discrodant PD-L1 expression is displayed as final scoring performed on full sections (marked with an asterisk). UPS, Undifferentiated pleomorphic sarcoma; Gde, FNLCC Grade (Fédération Nationale des Centres de Lutte Contre Le Cancer); Rec, Recurrence; Mets, Metastasis; TILs, Tumour-infiltrating lymphocytes.

| Case No. | Label ID. | Patient's age | Diagnosis | Site | Sample type | Year of Dx. | PD-L1 IHC % | TILs |
| --- | --- | --- | --- | --- | --- | --- | --- | --- |
| 23 | LM028 | 64 | Epithelioid leiomyosarcoma, 95mm | Uterus | Primary | 12 | 0 | No TILs |
|  | LM024 |  |  | Vagina | Mets | 13 | 0 | No TILs |
| 24 | LM015 | 60 | Log-grade leiomyosarcoma, 124 mm | Colon | Mets 1 | 12 | 0 | No TILs |
|  | LM119 |  |  | Colon | Mets 2 | 14 | 0 | No TILs |
|  | LM080 |  |  | Colon/bladder | Mets 3 | 16 | 0 | No TILs |
| 25 | LM099 | 48 | Leiomyosarcoma (Hx primary uterine) high grade, 24 mm | Lung | Mets 1 | 17 | *10% | High |
|  | LM100 |  |  | Lung | Mets 2 | 17 | *5% | High |
| 26 | LM020 | 56 | Leiomyosarcoma, high gde, 190mm | Uterus | Primary | 13 | *1% | No TILs |
|  | LM117 |  |  | Lung | Mets | 14 | *90% | No TILs |
| 27 | LM120 | 61 | Pleomorphic leiomyosarcoma, high gde, 50 mm | Abdomen | Primary | 14 | 0 | No TILs |
|  | LM106 |  |  | Lung | Mets | 17 | 0 | No TILs |
| 28 | LM062 | 76 | High grade leiomyosarcoma, 70 mm | Thigh | Primary | 08 | 0 | No TILs |
|  | LM056 |  |  | Thigh | Rec | 09 | 0 | No TILs |
| 29 | LM103 | 44 | Leiomyosarcoma (Hx primary uterine), high gde, | Pelvis | Rec | 17 | *1% | High |
|  | LM104 |  |  | Rectum | Mets | 17 | *5% | High |
| 30 | LM046 | 64 | Low gde epithelioid leiomyosarcoma, 15mm | Uterus | Primary | 10 | 0 | No TILs |
|  | LM030 |  |  | Omentum | Mets | 11 | 0 | No TILs |
| 31 | LM051 | 32 | Leiomyosarcoma, low grade, 124 mm | Uterus | Primary | 10 | 0 | Mild |
|  | LM116 |  |  | Lung | Mets | 15 | 0 | No TILs |
| 32 | LM019 | 43 | Leiomyosarcoma, primary, 110mm, gde 2 | Kidney | Primary | 13 | *5% | No TILs |
|  | LM118 |  |  | Omentum | Mets | 14 | *1% | No TILs |
| 33 | LM078 | 50 | Metastatic leiomyosarcoma (Hx primary uterine) | Lung | Mets 1 | 16 | 0 | No TILs |
|  | LM079 |  |  | Lung | Mets 2 | 16 | 0 | No TILs |
|  | LM085 |  |  | Lung | Mets 3 | 17 | 0 | No TILs |
| 34 | ML03 | 53 | Myxoid liposarcoma, 90 mm (poorly diff areas 5%) | Popliteal fossa | Primary | 15 | 0 | No TILs |
|  | ML15 |  |  | Thigh | Rec | 16 | 0 | No TILs |
| 35 | ML05 | 54 | Myxoid liposarcoma, 130 mm, low grade | Leg | Primary | 14 | 0 | No TILs |
|  | ML16 |  |  | Lung | Mets | 16 | 0 | No TILs |
| 36 | PLE-1 | 81 | Pleomorphic liposarcoma, Gde 3, 70 mm | Arm | Primary | 15 | 0 | No TILs |
|  | PLE-7 |  |  | Arm | Rec | 18 | 0 | No TILs |
| 37 | PLE-3 | 51 | Pleomorphic liposarcoma, post-irradiation, gde 1, 20 | Deltoid | Rec 1 | 13 | 0 | No TILs |
|  | PLE-5 |  |  | Deltoid | Rec 2 | 14 | 0 | Moderate |
| 38 | DD-LP28 | 64 | Dedifferentiated liposarcoma, High gde, 225 | Small bowel | Rec 1 | 8 | 0 | Mild |
|  | DD-LP26 |  |  | Retroperitoneum | Rec 2 | 9 | 0 | No TILs |
| 39 | AS-14 | 75 | Angiosarcoma, low and high grade, 325mm | Breast | Primary | 15 | 0 | No TILs |
|  | AS-15 |  |  | Chest wall | Rec | 15 | 0 | No TILs |
| 40 | AS-11 | 47 | Angiosarcoma, low grade, 50 mm | Atrium | Primary | 14 | 0 | No TILs |
|  | AS-01 |  |  | Atrium | Rec | 15 | 0 | No TILs |
| 41 | AS-08 | 79 | Angiosarcoma, high grade, 55m. Previous AS, 10 years | Vagina | Mets 1 | 12 | 0 | No TILs |
|  | AS-09 |  |  | Gingiva | Mets 2 | 12 | 0 | No TILs |
| 42 | RMS-01 | 57 | Pleomorphic rhabdomyosarcoma, 30 | Spermatic cord | Primary | 16 | *40% | High |
|  | RMS-08 |  |  | Brain | Mets | 18 | *80% | No TILs |
| 43 | RMS-17 | 17 | Embryonal rhabdomyosarcoma, 65 | Paratesticular | Primary | 14 | 0 | No TILs |
|  | RMS-18 |  |  | Pelvic LN | Mets | 14 | 0 | No TILs |
| 44 | SS1-B | 31 | Synovial sarcoma, 150 mm, SS18+ | Femur | Primary | 12 | 0 | No TILs |
|  | SS1-C |  |  | Lung | Mets 1 | 13 | 0 | No TILs |
|  | SS1-D |  |  | Lung | Mets 2 | 13 | 0 | No TILs |
|  | SS1-E |  |  | Thigh | Rec | 14 | 0 | No TILs |
|  | SS1-F |  |  | Lung | Mets 3 | 15 | 0 | No TILs |
|  | SS1-G |  |  | Lung | Mets 4 | 15 | 0 | No TILs |
| 45 | SS7-A | 70 | Synovial sarcoma, 55mm, SS18+ | Foot | Primary | 17 | 0 | No TILs |
|  | SS7-B |  |  | Leg | Rec | 17 | 0 | No TILs |
| 46 | RC-12 | 30 | Ewing sarcoma, post chemotherapy, 80 mm, EWSR1+ | Rib | Rec 1 | 14 | 0 | No TILs |
|  | RC-02 |  |  | Vertebra | Rec 2 | 17 | 0 | No TILs |
|  | RC-01 |  |  | Lung | Mets | 18 | 0 | No TILs |

Nineteen per cent of matched samples (9 of 47 patients) displayed true differences in the proportion of cells expressing PD-L1 in the primary tumour when compared to its corresponding metastasis / recurrence or between metachronous metastases / recurrent episodes. These include samples from 4 patients with MFS, 4 with LMS and 1 with pleomorphic RMS. In these samples, the percentage of PD-L1+ cells increased in the metastatic/recurrent site compared to the primary tumour in 6 of 9 cases (67%), from which 3 cases had only a modest increase (i.e. 1% to 5 or to 10%; **Table 1**) but a dramatic increase was identified in the 3 remaining cases (**Figure 2**). These included a uterine leiomyosarcoma with original PD-L1 expression of 1%, which increased to 90% in its lung metastasis (**Figure 2A-D**), a pleomorphic rhabdomyosarcoma from the spermatic cord with increase from 40% to 80% in the brain metastasis (**Figure 2I-L**) and a high-grade MFS with increase from 5% to 70% in the recurrence (**Figure 2E-H**). Interestingly, for this MFS case, gain in PD-L1 was identified in the second recurrence (Rec 2) with the first recurrence (Rec 1) only showing 1% expression (**Table 1**). PD-L1 was negative in all paired samples from angiosarcomas, myxoid liposarcomas, synovial sarcomas, pleomorphic liposarcomas and Ewing sarcomas.

**Figure 2 legend.**
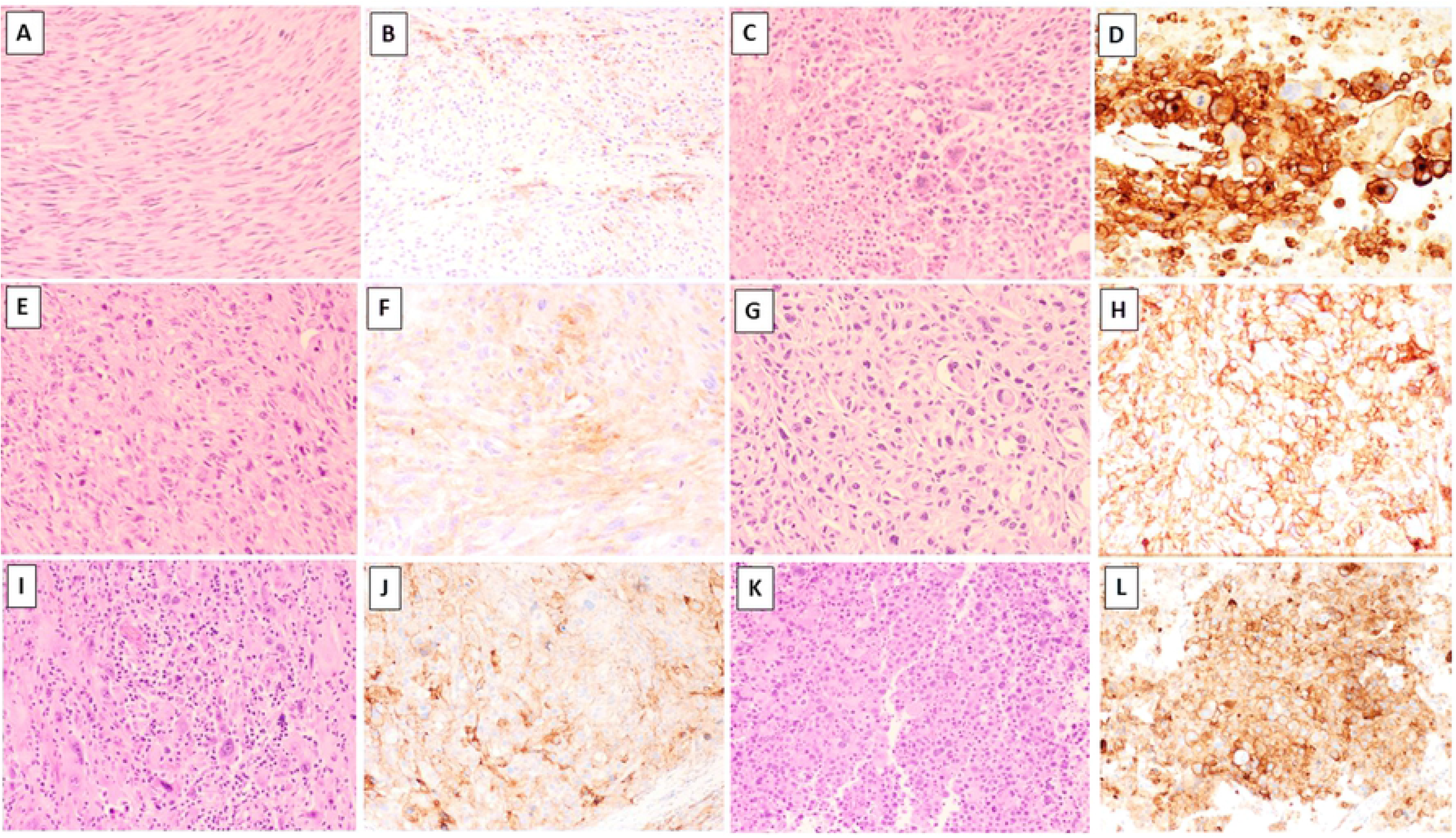
Matched samples (A-D, E-H and I-L, respectively). Primary uterine leiomyosarcoma diagnosed in 2013 (A, LM20) showing 1% weak expression for PD-L1 in the primary tumour (B). Lung metastasis of this tumour presented one year after the primary diagnosis (C; LM117), displaying strong PD-L1 expression in 90% of the tumour cells (D). High-grade myxofibrosarcoma (E, MFS-92) with weak PD-L1 expression in 5% of the cells (F). Recurrence one year later (G; MFS-72) demonstrated moderate to strong PD-L1 expression 70% of the tumour cells (H). Primary rhabdomyosarcoma (I; RMS-1) with 40% heterogeneous expression for PD-L1 in the original tumour (J), which increased to 80% in the brain metastasis (K-L). In this case it can also be noted the presence of tumour-infiltrating lymphocytes (TILs) in the primary tumour (1I) but not in the brain metastasis (1K). Sections stained with Haematoxylin and Eosin (A,C,E,G,I,K) or with the PD-L1 SP263 clone, Ventana (B, D, F, H, J, L). Images taken at 10X-40X magnification.

### TIL correlations

Correlation between PD-L1 expression and the presence and degree of TILs was investigated on full H&E sections exclusively on the sarcoma subtypes with the highest number of cases (MFS, UPS and LMS) and for the Mets/Rec cohort **(Table 1**). MFSs and UPSs were combined for the analysis as it has been shown that these appear to represent one single biological entity within a phenotypic spectrum(21). Concordant TIL and PD-L1 expression on the same cases was available for 158 UPS/MFS cases, which showed TILs in 41% of all cases (n=64) with 17%, 17% and 8% showing minimal, intermediate and high TILs (**Figure 3**), respectively. No association was identified with overall TIL levels and PD-L1 expression (P value 0.06 and Z-score −1.8; Mann Whitney test). To investigate further any possible correlation, TIL groups were dichotomised as cases with absent-to-minimal (<10%) TILs vs. those with moderate-to-marked TILs (>10%). UPS/MFSs with minimal-to-absent TILs expressed PD-L1 in 22% of cases (n=26) and those with moderate-to-marked TILs expressed PD-L1 in 31%. This was once again, not statistically significant (P= 0.28, Fisher’s exact test).

**Figure 3 legend.**
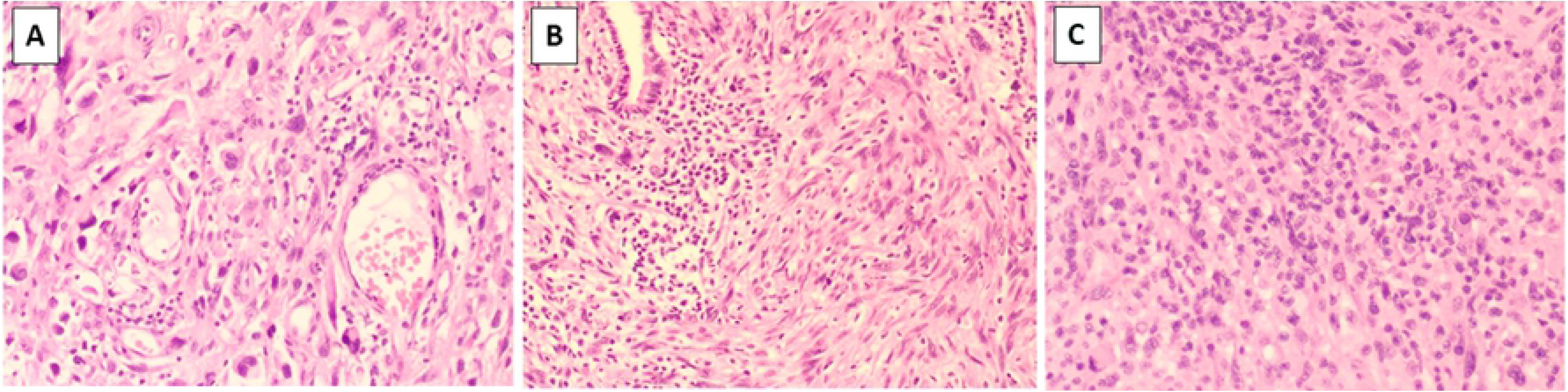
Stromal Tumour-Infiltrating Lymphocytes (TILs) on sections stained with Haematoxylin and Eosin (H&E). The images show examples of degree of TILs classified as nil (0%, not shown), minimal (1-10%, A), intermediate (>10% and <50%, B) and high (>50%, C).

For the leiomyosarcoma group (n=116), TILs were rare overall with 83% (n=96) of all cases showing complete absence of TILs and with the remaining 17% having any degree of TILs (8%, 4% and 6% with minimal, intermediate and high TILs, respectively). Once again, no significant correlation was identified between overall PD-L1 expression and presence or absence of TILs in LMSs (P=0.5; Z-score 0.66, Mann Whitney test). When the cases were dichotomised, it was observed that PD-L1 expression was present in 5% (5 of 105) of cases with minimal-to-absent TILs vs. 72% (8 of 11) in those LMS with moderate-to-marked TILs (8 of 11: 72%; <0.0001, Fisher’s exact test).

In the Met/Rec cohort, no correlation was identified between PD-L1 expression and TILs including those cases associated with apparent gain of PD-L1 in the metastatic or recurrent deposit (**Table 1**). An inverse correlation was identified in one of the three cases with significantly higher PD-L1 expression in the metastasis (pleomorphic RMS), as this showed absent TILs in the brain metastasis but high expression in the primary tumour, whereas PD-L1 expression was higher in the metastasis compared to the primary tumour (80% vs. 40%, respectively; **Figure 2I-L**). The other paired cases showed no TILs at initial presentation and remained absent in the metastatic/recurrent disease (**Table 1**). Lastly, in this Met/Rec cohort, changes in the histomorphological features were also assessed between primary tumours and their matched recurrent/metastatic samples on full H&E sections to determine whether gain in PD-L1 and the metastatic/recurrent site was associated with changes in the morphology of the primary tumour. Although some samples showed changes, a significant trend was not observed.

## Discussion

There are several well-described factors which account for the variability in the reported expression of PD-L1 in lung adenocarcinomas and sarcomas (1, 18, 28-32), making comparisons of the literature challenging. One of these variables is the use of over 10 commercial and non-commercial PD-L1 antibodies including SP263 and SP142 (Ventana Medical Systems), E1L3N (Cell Signalling Technology), 22C3 (PharmDx kit, Agilent Technologies), 28-8 (PharmDx kit), 5H1 (L. Chen, John Hopkins University), #ab58810 and EPR1161(Abcam) and #SAB2900365 (Sigma-Aldrich). Clones differ in the binding site to the PD-L1 protein: extracellular (28-8 and 22C3, SP263 and E1) vs. cytoplasmic (SP142 and E1L3N) (Reviewed in(1)) with further technical differences regarding antigen retrieval conditions, staining platforms and differential binding properties according to cell types (i.e. epithelial vs. immune). The use of inconsistent cut-offs is also a significant contributing factor to discordant PD-L1 immunohistochemical expression, and it has been shown that by adjusting predefined cut-offs this can lead to misclassification of PD-L1 status in tumour samples (30). Finally, biological reasons also contribute to the lack of uniformity in PD-L1 expression. For instance, PD-L1 can be constitutive or inducible in response to IFN-γ released by effector T cells. Inducible PD-L1 can be affected by tumour location and prior therapy (Reviewed in(1)) including radiotherapy (33). At the current time, there is not enough literature to address the dynamics of inducible PD-L1, as a result of therapy but if such changes occur, these may potentially provide therapeutic avenues to patients who may not initially be suitable for anti-PD-L1 therapy.

Although providing an average incidence of PD-L1 expression in our combined sarcoma cohort is not biologically relevant, in view of the different molecular signature of soft tissue sarcomas(20, 23), PD-L1 was present in 13% of all cases in which was the protein essentially only identified in tumour cells and not in immune cells when assessed primarily on TMAs. Our study shows relatively concordant findings to those previously published with the SP263 clone on smaller number of selected sarcoma subtypes(10, 18). In this study, we demonstrated the highest expression of PD-L1 in UPS (31%), followed by angiosarcomas (29%), RMS (26%, predominantly pleomorphic subtype), MFS (18%), LMS (11%) and DD-LPS (10%). Sarcomas uniformly negative for PD-L1 expression in our series included ALT/WDL, myxoid liposarcomas, synovial sarcomas, pleomorphic liposarcomas, and Ewing sarcomas. This is in line with the previous observation that PD-L1 expression is a feature of sarcomas with a complex karyotype with or without high mutation burden (18) and only seen in our cases related to the complex karyotype group (UPS, angiosarcomas, pleomorphic RMS, MFS, LMS and DD-LPS). Our results however, differ when compared with other similar studies using antibodies other than the SP263 and variable cut-offs (i.e. >2+ and >5%(12)). In some of these series, the frequency of PD-L1 expression ranges from absent (0%) with the Abcam (ab58810 and ab205921) and SP142 clones(7, 8) to low (12%: H-130 clone)(9), intermediate (26-30%: AM2653AF-N. Acris & SP263)(10, 11) and high (58% & 65% with BD Pharmingen and R&D Systems and the H-130, Santa Cruz clones, respectively(12, 13)). Although strong correlation has been established for some clones (i.e. SP263 and 22C3) but less for others (SP142(18)), such comparative analyses have not been widely performed in sarcomas as opposed to lung adenocarcinomas. Our results are also discordant with studies reporting high PD-L1 expression in some specific sarcoma entities not identified in our cohort (i.e. liposarcomas(16), myxoid liposarcomas(12), synovial sarcomas(13) and Ewing’s sarcoma family of tumours (13, 34).

We did not investigate the correlation between PD-L1 IHC expression and specific clinical pathological variables (i.e. tumour size, grade, site, patient’s age, etc), as such correlation has not been previously identified (9, 14) and no obvious trend was seen in our cohort (See Table 1 for clinical pathological details shown for the Met/Rec cohort). In the largest meta-analysis published to date(14) including 1451 sarcoma patients, Zheng et al, suggested that PD-L1 expression does not correlate with patient age, histological grade, clinical stage, site, necrosis, chemotherapy, radiotherapy or incidence of metastasis or recurrence. This study, however, confirmed other studies showing that PDL-L1 expression is a poor prognostic factor for event-free survival in both soft tissue sarcomas and osteosarcomas. A significant limitation of the Zheng et al meta-analysis is that sarcoma subtypes were combined for statistical analysis, no taking into account the biological diversity of the different types of sarcomas. Therefore, large studies addressing the clinical diversity of soft tissue sarcomas stratified based on PD-L1 expression are lacking. Clinical follow-up is not available in our cohort.

With regards to increase / gain of PD-L1 in the metastatic setting, limited studies have documented similar findings in different tumour types. Lussier et al, in a study of 11 paediatric osteosarcomas, demonstrated that 75% of metastatic lesions but no primary osteosarcomas expressed PD-L1(35). The same phenomenon was observed in a case of dedifferentiated chondrosarcoma, which showed absence of PD-L1 in the primary tumour and positive expression in the metastasis(36) and this has also recently been documented in a series of metastatic renal cell carcinomas(37). Here, we report for the first time that gain in PD-L1 protein expression can occur in the context of metastatic/recurrent myxofibrosarcoma, leiomyosarcoma and pleomorphic rhabdomyosarcoma. We demonstrated on full section analysis that the percentage of positive cells expressing PD-L1 increased in the metastatic/recurrent site compared to the primary tumour in a total of 6 of 9 cases with a particularly marked increase in 3 of those cases. Based on results of the KENOTE-010 study, patients with high PD-L1 score (>50%) have significantly increased benefit with pembrolizumab compared to those with >1%(3). Therefore, it is possible that these patients, all of which showed >50% positive cells in the metastasis/recurrence but not in the primary tumour, may have an objective treatment response. Contrary to these findings, the few comparative similar analyses between paired primary and metastatic lung carcinoma samples have identified that discrepant PD-L1 expression, when it occurs, is due to negative expression in the metastatic deposit/s compared to primary tumours (8, 27, 38, 39). It should be noted that in some of these studies, biopsies rather than resections have been used to determine concordance and intra-tumour heterogeneity for PD-L1 has been shown to account for the apparent discordance in some of those series.

With regards to our analysis of TILs assessed on H&E sections in a subset of cases, we identified no correlation with PD-L1 expression in the Met/Rec cohort but a significant association was identified for a specific sarcoma subset, leiomyosarcomas (n=116, full subset). None of the three cases of the Met/Rec cohort with significant increase in the percentage of PD-L1 positive cells contained TILs. In fact, one of those cases had high TILs in the primary and absence in the brain metastasis (**Figure 2I-L**). Lack of TILs in brain metastases has been previously reported(38). Our other Rec/Met paired cases showed concordant absence of TILs at initial presentation and on the metastasis / recurrence (**Table 1**). As high density TILs (specifically, CD3+/CD8+) is an independent positive prognostic factor for OS and DFS in sarcomas(10), it would be expected that absence of TILs would correlate with aggressive behaviour in those sarcomas prone to recur and/or metastasise. Induction or oncogeneic activation of PD-L1 in non ‘inflamed’ tumours occurs through alternative pathways such as loss of PTEN or copy gain / amplification of CD274/PDL1 locus (Ch 9p24.1)(10, 15). In our full cohort of leiomyosarcomas (n=116), PD-L1 expression strongly correlated with the presence of moderate-to-high TILs identified on H&E sections when compared to absent-to-minimal (<10%) TILs (<0.0001, Fisher’s exact test). In a recent series of uterine smooth muscle tumours including 23 LMSs (40), Shane et al, demonstrated higher CD3+ TILs in leiomyosarcomas but direct correlation with co-expression of PD-L1 was not assessed(40).

We did not characterise subsets of TILs (ie.. CD8+, FoXP3+, etc) because this has been previously published(5, 7–19) and because our study focused on the possible relevance of TILs on histological H&E-stained sections. With regards to the correlation between PD-L1 and PD-1 expression this is conflicting with a study from the Cancer Genome Atlas Research Network demonstrating that PD-L1 mRNA does not correlate with PD-1(21). This has also been documented in smaller studies (7).

Our study has a number of limitations. First of all, it is essential to emphasise that the clinical predictive value of PD-L1 protein expression assessed on tumour samples has not been widely investigated in sarcoma patients and there is currently only scanty evidence from single-arm phase 2 studies of response to PD-L1 inhibitors. Importantly, in the SARC028 trial clinical, response to checkpoint inhibitors was seen in cases even in the absence of PD-L1 expression (5), which may indicate that PD-L1 expression may not necessarily impact treatment response. Nonetheless, 2 of the 3 patients with positive PD-L1 expression in that trial (4% in total; 3 of 70 patients with a range of sarcoma types) showed either complete or partial treatment response. This is suggestive of some predictive value of PD-L1 expression assessed in pre-treatment biopsies (5). PD-L1 expression on the full cohort was assessed using TMAs. This was partly overcome by performing PD-L1 IHC and assessment of TILs on full sections. There is no clinical follow-up available for our cases including a lack of details about treatment modality in the vast majority. Importantly, the cohort predominantly includes treatment-naïve sarcomas with the exception for the Met/Rec cohort. For this subset of cases, it is expected that patients were treated with conventional regimens according to sarcoma type. Hence, it is likely that they were treated following universal guidelines before clinical trials were more widely available and according to clinical stage. This would make comparative analyses reliable. Although radiotherapy has been shown to induce PD-L1 expression(33), our cases with documented post-radiotherapy (Table 1; n=5) showed <10% PD-L1 expression. Finally, the size of the metastatic/recurrent cohort is limited and therefore a larger number of paired samples is needed to further validate these findings. Nonetheless, paired samples are scant in most centres but comprise very valuable resource to understand the biology of sarcoma progression.

## Conclusions

Overall, we conclude that the frequency of PD-L1 expression in sarcomas is limited to some sarcoma subtypes and that variable selection of clones and cut-offs contributes to the marked discrepancy across different series. We focused on the 263 companion kit as this selectively identifies patient’s eligibility to anti-PD-1 and/ PD-L1 inhibitors in some other cancers. Our study showed that PD-L1 can objectively increase in a small proportion of cases in paired metastases/recurrent disease to >50% expression in the tumour cells. This suggests the possibility of treatment response to immune checkpoint inhibitors in the metastatic/recurrent setting but unlikely in the primary tumour in a proportion of cases (i.e. as PD-L1 of 1-5%). In our assessment of TILs on H&E sections, we demonstrated that PD-L1 expression in leiomyosarcomas was associated with at least moderate levels of TILs identified on H&E sections. Although the prognostic significance of this association cannot be elucidated in this study, in view of the lack of clinical follow-up, the positive correlation of TILs and PD-L1 may help to triage the LMS cases which may benefit from PD-L1 IHC. Our results indicate that LMSs with moderate TILs are significantly more likely to harbour PD-L1 expression.

## Acknowledgements

We thank A/Prof Annabelle Farnsworth and Amy Sheen, who assisted with ethics approval. Andrew Townsend, Leanne Henderson and Natalie Jukic collaborated with database search and tissue retrieval of the sarcoma cohort at Douglass Hanly Moir (DHM) Pathology. We also thank the histopathologists from DHM Pathology who reported a proportion of the cases presented in this study: Prof Warick Delprado, Dr Ivan Burchett, Dr Julie Burn, Dr Tina Baillie, Dr Debra Jenson, Dr Cate Trebeck, Dr Robyn Levingston, Dr Patricia Guzman, Dr Claire Biro, Dr Helen Ogle, Dr Elizabeth Sinclair, Dr Geoffrey Hall, Dr Alexandra Allende, Dr Suzanne Hyne, Dr Anita Muljono, Dr Jennifer Roberts, A/Prof Jennifer Turner, Dr Melanie Edwards, Dr Denis Moir and Dr Martin Jones. We also acknowledged the contribution from external histopathologists from RPA Hospital: Dr Rooshdiya Karim and Prof Stan McCarthy.

